# Predictive adequacy and measurement dependence of metabolic representations in controlled synthetic systems

**DOI:** 10.64898/2026.10.27.762156

**Authors:** Mwangi Njenga, Waweru Peter

## Abstract

**Background:** A metabolic representation may predict adequately, appear compact or change repeatably without satisfying the other properties. Evaluating these properties requires explicit reference information, measurement conditions and definitions of change.

**Methods:** Six stipulated stochastic scenarios generated twenty independent confirmation cohorts each, with 5,115 synthetic individuals per cohort and 1,023 held out. Three representation families—selected current and longitudinal descriptors (Φ), a larger selected collection of measurements and historical descriptors, and an outcome-guided learned embedding of longitudinal inputs—were evaluated against complete available longitudinal history for year-25 prediction from information available through year 20. Forty-two adequacy groups used overall and context-specific relative-loss margins with 2,850 simultaneous upper bounds. Information accounting distinguished displayed coordinates, upstream values and nominal input fields. Separate procedures examined current remeasurement, natural coordinate tracking and controlled endpoint responses.

**Results:** Five groups satisfied every adequacy requirement; none of the twelve Φ groups did. Of 393 failed predicates, 321 had estimates within the margin but upper bounds above it. The conventional-panel embedding displayed eight coordinates but depended on 72 upstream engineered values and additional baseline, context, support and missingness inputs. Remeasurement stability did not ensure lower prediction error, and changing the future scoring reference could reverse a method ordering without changing predictions. Natural tracking yielded 25,440 defined associations, including 442 negative values, and 1,440 undefined associations. Controlled responses agreed with their assigned directions when numerically resolved, but included mixed dose sequences and unsupported coordinates. Of 47,127,708 eligible stochastic response records, 86.331% were smaller than one matched measurement standard deviation; 96.587% of records exceeding that reference retained their corresponding category.

**Conclusions:** Adequacy, compactness, repeatability and responsiveness describe different properties of a representation. Non-certification does not necessarily demonstrate excess loss, and category-silent movement does not establish clinical importance or subsequent predictive harm. Evaluation should retain complete input requirements, measurement references and unsupported cases. These synthetic findings do not establish a sufficient human physiological representation or therapeutic benefit.

## Introduction

A metabolic representation is useful only in relation to what it is expected to do. A few displayed coordinates may support prediction, but their construction can require many earlier observations. A stable value can reflect a stable underlying condition, insensitivity to change or an encoding that rarely crosses a boundary. Conversely, movement can arise from evolving physiology, measurement variation or both. These distinctions matter when representations are proposed for longitudinal assessment, where apparent simplicity and responsiveness can encourage interpretations that their evaluation has not established.

Continuous metabolic severity scores and longitudinal models provide substantive precedents. Factor-based severity scores summarise covariance among conventional measurements, and subsequent studies have related severity and its evolution to human outcomes (Gurka et al., 2014; DeBoer et al., 2015; Vishnu et al., 2015). Joint trajectory models represent correlated changes over time (Breeze et al., 2016). These approaches demonstrate the value of alternatives to a single categorical assessment. They do not make every compact score sufficient for every future outcome, nor establish that a change has the same meaning under different measurement conditions.

Measurement invariance provides a related caution. Stable relationships between observed indicators and a statistical construct cannot simply be presumed throughout follow-up (Barbosa-Leiker et al., 2011). In a dynamical setting, history may also improve estimation of the present when current measurements are noisy, even when the underlying state follows a Markov law (Särkkä & Svensson, 2023). Predictive adequacy must therefore identify its reference information and task, while responsiveness must identify the source and scale of the change being examined.

We studied these questions within controlled synthetic metabolic systems. The systems separated latent physiology, its observations, the record available at a prediction cutoff and the representation supplied to a learner. This permitted four connected assessments: whether specified representations met a declared predictive requirement relative to available history; what information their construction consumed; how repeated current measurements affected their outputs and predictions; and how their coordinates responded to natural evolution or a controlled input change. The objective was an assessment of distinct properties, without assuming that Φ or any other representation should satisfy all of them. Simulation supplied experimental control conditional on stipulated laws, rather than an estimate of human validity (Morris et al., 2019).

## Methods

### Study design and information boundaries

Six stochastic scenarios represented current-state-sufficient dynamics, accumulated burden, reserve depletion and recovery, an adapting operating reference, persistent historical effects, and nonlinear regime-dependent dynamics. Four interacting domains represented adiposity, glycaemia, lipoprotein metabolism and haemodynamics. Scenario-specific latent quantities influenced subsequent transitions and generated measurements. They were mathematical constructs with declared roles, not identified human physiological compartments. Generating parameters, initial distributions and observation mechanisms were stipulated rather than fitted to human cohorts. Supplementary Methods S1 specifies the limited study generator; the observation and availability rules are given in Supplementary Methods S2.

Four development cohorts of 1,024 individuals per scenario supported recipe selection. Confirmation used twenty independent cohorts of 5,115 individuals per scenario, partitioned into 3,069 training, 1,023 validation and 1,023 test individuals. All records and future branches for one individual remained in one partition. The same 120 confirmation cohorts supported repeated observation and panel conditions; these repeated views did not create additional independent cohorts. Three principal panels contained six, nineteen and thirty-five measurements. The conventional panel comprised waist circumference, systolic and diastolic blood pressure, triglycerides, HDL cholesterol and fasting glucose.

Four observation conditions supplied complete noiseless annual observations, scheduled noisy observations, less-frequent scheduled observations, or a joint observation–treatment-policy stress condition. The noisy settings included persistent individual–channel, shared-visit and observation-specific variation. The joint stress setting also changed missingness, reporting delay and treatment-influenced physiology, so it did not isolate measurement noise. Predictor eligibility required acquisition and availability by the queried time, with the prespecified age and historical-gap restrictions. Missing measurements were not replaced by access to latent states.

Information through year 20 supported prediction of six conventional measurements at year 25. Future branches shared earlier history and paired process innovations. In one branch external loading continued; in the other each controlled input was set to 0.05 at year 20. We call the latter the fixed-low-load branch, corresponding to the historical withdrawal label. Setting a fixed input value was not guaranteed to reduce every individual’s input and did not remove internal burden. Both branches were synthetic regimes, not clinical treatments. The complete design produced 1,440 main observation–panel cells; forty restricted research-extension cells were separate settings.

### Representations and fitted prediction

The principal candidates were Φ, an explicit collection of selected current and longitudinal descriptors; a larger selected collection of measurements and historical descriptors; and an outcome-guided learned embedding of longitudinal inputs. Their recipes and dimensions were fixed using development data before confirmation. Φ retained eight, sixteen or thirty-two value features across the three panels. The learned embedding retained eight, twelve or twelve coordinates, obtained through a supervised linear projection. Selection or compression did not establish physiological identity, minimum dimension or adequacy.

Complete Available Longitudinal Measurement History was the predictive reference. It contained eligible scheduled measurements and its declared auxiliary fields, with no privileged access to latent physiology. It was a fitted reference, not a Bayes-optimal oracle. Baseline measurements, context, support fields and missingness indicators accompanied value blocks where specified. Current measurements and hard clusters were included in the remeasurement comparisons. Cluster labels used frozen fitted maps and were encoded without an assumed severity ordering.

Ridge regression used an unpenalised intercept and nine candidate penalties from 10⁻⁴ to 10,000. Validation selected the penalty by equally weighted, scale-normalised outcome loss separately for each task; the selected configuration was refitted using training plus validation. Preprocessing, feature selection and projection estimation respected their prescribed fitting partitions. Confirmation test outcomes did not choose recipes or tune models. The embedding teacher used a fixed Ridge penalty of one. The relevant definitions and support rules remained unchanged across the analyses reported here. Supplementary Methods S3 describes the representation definitions, locked Φ coordinates and fitting boundaries.

### Prediction loss and adequacy

Channel-specific loss was the held-out mean squared prediction error divided by the squared standard deviation of the corresponding observed training outcome. That between-individual normalising scale was distinct from measurement error. The six-channel endpoint weighted channels equally; these were experimental weights rather than clinical-importance weights. Main scoring used complete synthetic noisy year-25 outcomes, whereas fitting targets retained their specified missingness. Noiseless future references belonged to a separate descriptive analysis.

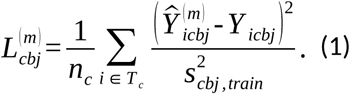

The overall assessment used twenty prespecified replicate-index blocks, each containing the confirmation cohort with the same index from each of the six scenarios. Within each block, candidate losses and available-history reference losses were separately averaged with equal weights over the six scenarios, two future tasks and six channels. Relative excess was then calculated from these two averages, and the twenty resulting ratios were averaged. Each scenario–task–channel guard instead used the twenty cohort ratios for its fixed scenario, task and channel. Specialist overall means covered the thirty-six scenario–channel cells for their assigned task. Supplementary Methods S4.3 gives the formulas and exceptional-case rules. Thirty-six common groups, defined by representation, panel and observation condition, jointly covered both future tasks. Each required one balanced overall predicate and seventy-two scenario–task–channel guards. Six specialist groups covered their assigned task under scheduled noisy observation, each with one overall predicate and thirty-six scenario– channel guards.

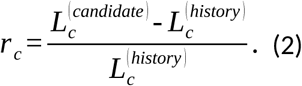

A predicate’s upper confidence bound was its mean relative excess plus its standard error multiplied by the Student t quantile with nineteen degrees of freedom at 1−0.05/2,850. For every predicate, the standard error was the sample standard deviation of its twenty replicate-level quantities, calculated with divisor nineteen, divided by the square root of twenty. Thus, the simultaneous family contained 36×73+6×37=2,850 one-sided predicates. Certification required an overall upper bound no greater than 2% and every applicable guard no greater than 10%. These margins were operational choices for this synthetic study, not clinical non-inferiority margins. The t reference is exact for independent, identically distributed normal cohort summaries and otherwise approximate; the Bonferroni allocation preserves its simultaneous interpretation only to the extent that the individual bounds have their stated coverage.

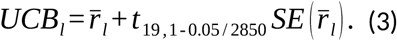

A failed requirement meant that adequacy was not certified. An upper bound above a margin did not necessarily show that the true excess exceeded that margin, and failure of a two-task conjunction did not establish failure separately in both tasks. The certificate also differed from formal predictive sufficiency, which concerns conditional distributions rather than the loss of a particular fitted reference (Dawid, 1979). Undefined evidence retained its prescribed status. Splitting the report did not recalculate or reduce the original simultaneous family. Supplementary Methods S4 retains the full predicate structure and exceptional-case rules.

### Accounting for information requirements

We distinguished representation values, upstream engineered inputs, nominal input-field counts, distinct measurement channels, observation occasions and historical coverage. A learned coordinate could depend on several input features, and an input feature could depend on repeated observations of one channel. Baseline, contextual, support and missingness fields were counted separately from the displayed value block. Nominal input-field counts followed the accounting convention in Table 4 and excluded additional indicators introduced by downstream preprocessing. Realised usable nonconstant columns could vary between cohorts.

The channel counts associated with selected value blocks did not, by themselves, give the union of channels required by every appended input. In particular, the ten- and fourteen-channel counts for the larger-panel Φ blocks were not treated as verified reductions in total acquisition requirements. A complete union across all appended fields was not established by the available accounting. No monetary cost, patient burden, optimal measurement schedule or globally minimum physiological dimension was estimated.

### Fixed physiology with repeated current measurement

The remeasurement procedure used the first 256 held-out individuals in each of 120 conventional-panel, scheduled-noisy scenario–cohort contexts. Thirty current-observation replicas were generated while latent state, earlier observed history and fitted pipelines remained fixed. Prediction spread was the finite-replica variance around each individual’s replica-mean prediction, using divisor thirty. Errors against the saved noisy and noiseless year-25 references were calculated from the same predictions and retained the frozen channel-specific squared scale. Table 1 distinguishes this population from those of the other movement procedures. For a fixed individual, fitted method, future task and outcome channel, let *p_r_* denote the predicted year-25 measurement obtained from current-observation replica *r*, where *r* = 1 *,…,* 30. Predictions are expressed in the outcome channel’s native units. The quantity *p̅* is the mean prediction across those thirty replicas, and *V _R_* is their finite-replica variance, calculated with divisor thirty. Subject–channel bias and earlier observed history were held fixed, while a channel-shared visit error and channel-specific residual errors were redrawn for each current replica.

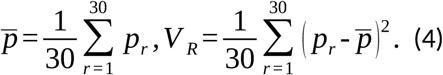

**Table 1.** Populations and endpoints of the three movement procedures.

| Procedure | Population | Endpoint | Reduction unit |
| --- | --- | --- | --- |
| Current remeasurement | 120 conventional-panel scheduled-noisy contexts; first 256 held-out individuals; 30 replicas | Prediction spread, future-reference error and clean-input displacement | Replicas within individual, then prescribed equal-cohort summaries |
| Natural tracking | 1,440 repeated observation–panel cells; 122,760 distinct cohort–subject identities | Coordinate change and Spearman association with latent-change magnitude | Finite pairs within each cell; descriptive cohort summaries |
| Controlled endpoint response | 120 conventional-panel scheduled-noisy contexts; 30,720 subject–cell identities | Endpoint displacement, direction, dose ordering, category change and measurement-relative magnitude | Repeated coordinate, input-domain, dose and applicable replica records |

These populations overlap. Repeated panels, observations, coordinates and doses do not increase the number of independent cohorts. Natural tracking uses 1,023 held-out individuals per cohort; the other two procedures use 256. Full definitions and eligibility rules are in Supplementary Methods S5.

The noiseless reference was the mapped value on one realised future biological trajectory. It was not an expectation over future innovations or a clinical gold standard. A separate clean-current-input anchor removed current measurement error from the same fitted pipeline while preserving earlier noisy history, physiology and fitted state. Its endpoint was the squared difference between the replica-mean prediction and this clean-input prediction. Prediction spread, future error and clean-input displacement therefore answered different questions. Subject-level quantities were reduced according to the existing hierarchy, with equal-cohort summaries across twenty cohorts. Let *y*_25*,noisy*_ and *y*_25*,noise*-_ *_free_* denote the individual’s saved noisy and noise-free year-25 references for the same future task. Both references remain fixed across the thirty current-observation replicas. The noise-free reference is the mapped measurement on the same realised future biological trajectory with observation error removed. Equation (5) gives the mean squared error of the thirty predictions against each reference; changing the scoring reference does not change the predictions.

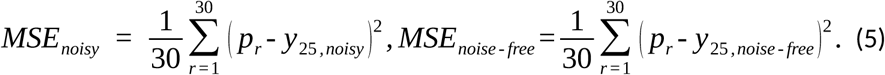

The mean prediction *p̅* retains the outcome’s native units. Prediction variance, both future-reference mean squared errors and clean-current-input squared displacement have native squared units. Their reported standardised values divide these squared quantities by 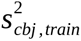, the frozen population variance of the original observed year-25 training outcomes for cohort *c*, future task *b* and channel *j*, calculated with divisor *n* rather than *n* - 1. The same denominator is used for both future references and is not re-estimated from the remeasurement replicas, noise-free outcomes or refitted prediction-head preprocessing.

Reconstruction examined where variation entered the representation pipeline. Raw-current preservation, fixed-history invariance and learned-coordinate variation were evaluated only where their inputs were defined. Reconstructed operations did not constitute independent observations. Learned coordinates remained specific to their fitted cell, and no common biological interpretation or universal stability metric was imposed on axes from different fits.

### Natural coordinate tracking

Φ was reconstructed at years 10 and 20 from information eligible at each respective time. The analysis used all 1,023 held-out individuals across 1,440 main cells, representing 122,760 distinct cohort–subject identities. Latent-change magnitude was the root mean square of changes in the four burden domains, each normalised by its designated training population’s year-20 standard deviation. This scoring quantity was not a predictor input or a clinical definition of meaningful change.

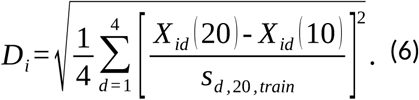

Within a cell, Spearman correlation related this latent-change magnitude to the absolute change in each eligible Φ coordinate. Signed coordinate changes were retained separately. Constant or insufficient pairs remained undefined. Cohort means and observed ranges were descriptive; no new significance tests or confidence intervals were introduced. No whole-vector Φ distance was established. Selected and learned constructions requiring observations after year 10, and research measurements requiring year-19 probes, could not supply lawful whole-vector year-10 comparisons. Their unavailable status was preserved.

### Controlled endpoint responsiveness

Starting from each saved year-15 state, one external-input domain was increased by 10%, 20% or 30% of its remaining headroom to 0.80, with other inputs unchanged. Zero-headroom cases remained included. The first 256 held-out individuals in each of 120 conventional-panel scheduled-noisy contexts were advanced monthly to year 20 and compared with matched unchanged-input controls. One lane used zero subsequent process innovations and clean endpoint values. A second lane used matched process innovations and three paired observation replicas.

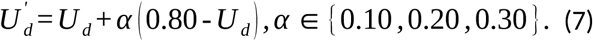

Both lanes preserved the original observed history through year 19 and substituted the new year-20 endpoint before applying the frozen representation. Consequently, this assessed endpoint responsiveness after a controlled continuation, not a fully re-observed counterfactual history. A future prediction head was not evaluated. Endpoints included displacement, agreement with predeclared directions, ordering across the three doses, and category change under frozen maps. No direction was invented after inspection for an unsupported regime or a learned axis.

Numerical resolution required absolute displacement to exceed twice the absolute tolerance plus the relative tolerance multiplied by the sum of absolute perturbed and control values. Smaller differences remained stored, rather than being converted to zero. In the stochastic lane, measurement-relative magnitude R was absolute coordinate displacement divided by the matched subject–coordinate remeasurement standard deviation. R required finite positive reference variance; zero and undefined references did not receive an arbitrary denominator. Category silence was summarised separately among resolved changes and among responses with R greater than one. These counts repeated individuals across coordinates, domains, doses and applicable replicas. Supplementary Methods S5 specifies the endpoint constructions, numerical tolerances and denominator rules.

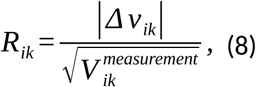

### Analytical chronology and reproduction limits

The analyses retained their original roles in the research programme. Development locks and confirmation adequacy procedures were distinguished from later readings of saved results, reconstruction, natural tracking and controlled-response releases. A plan fixed before a later calculation did not remove prior knowledge of related findings. The present organisation does not make these procedures one new prospective study. Descriptive endpoints were not assigned new p-values, and existing uncertainty or multiplicity rules were not changed for this report.

The historical predictive inventory contained 207,237 materialised results from 207,240 planned tasks; three remained unavailable. This inventory described computations across the wider programme, not the number of independent replications for any result here. Result inspection, reproduction of statistical summaries and reproduction of the entire experiment require different assets. Recorded identity and numerical-agreement checks supported bounded reconstruction, while incomplete historical dependencies and restart discrepancies limited claims of exact replay. The reporting record alone does not establish unrestricted experimental reproducibility. Supplementary Methods S7 distinguishes the study information disclosed with this report from execution assets and external result rosters that are not supplied.

## Results

### Adequacy depended on condition and task scope

Five of forty-two groups satisfied every requirement: four common groups and one specialist group (Figure 1; Tables 2 and 3). Thirty-seven were not certified; none had no verdict. None of Φ’s twelve common groups certified. Selected descriptors and the learned embedding each certified in two conventional-panel conditions, and no common group certified in either larger panel. These decisions jointly assessed their nominated tasks and contexts; they did not classify a representation as generally adequate or inadequate.

**Figure 1.**
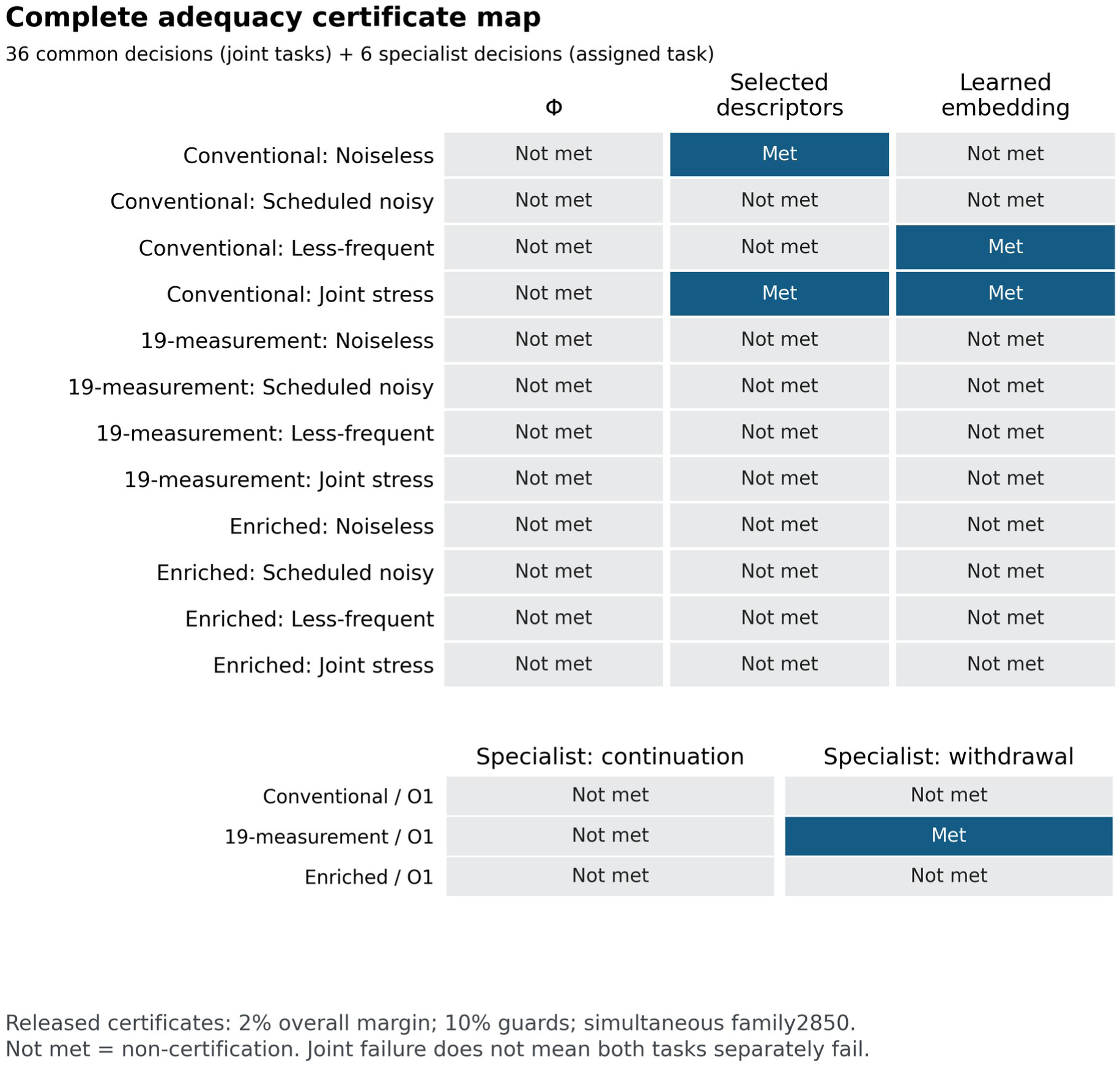
Adequacy depended on representation, observation and task scope. Five of forty-two groups met every simultaneous upper-bound requirement; none of the twelve Φ common groups did. Common groups jointly cover both future tasks and specialist groups cover their nominated task. The original 2,850-predicate family is retained. A failed joint requirement does not necessarily demonstrate excess loss beyond a margin or separate failure in both tasks. O0–O3 denote complete noiseless annual, scheduled noisy, less-frequent scheduled and joint observation–treatment stress conditions, respectively; P0–P2 contain six, nineteen and thirty-five measurements. Historical “withdrawal” labels, where shown, denote assignment to the specified low-load regime.

**Table 2.**
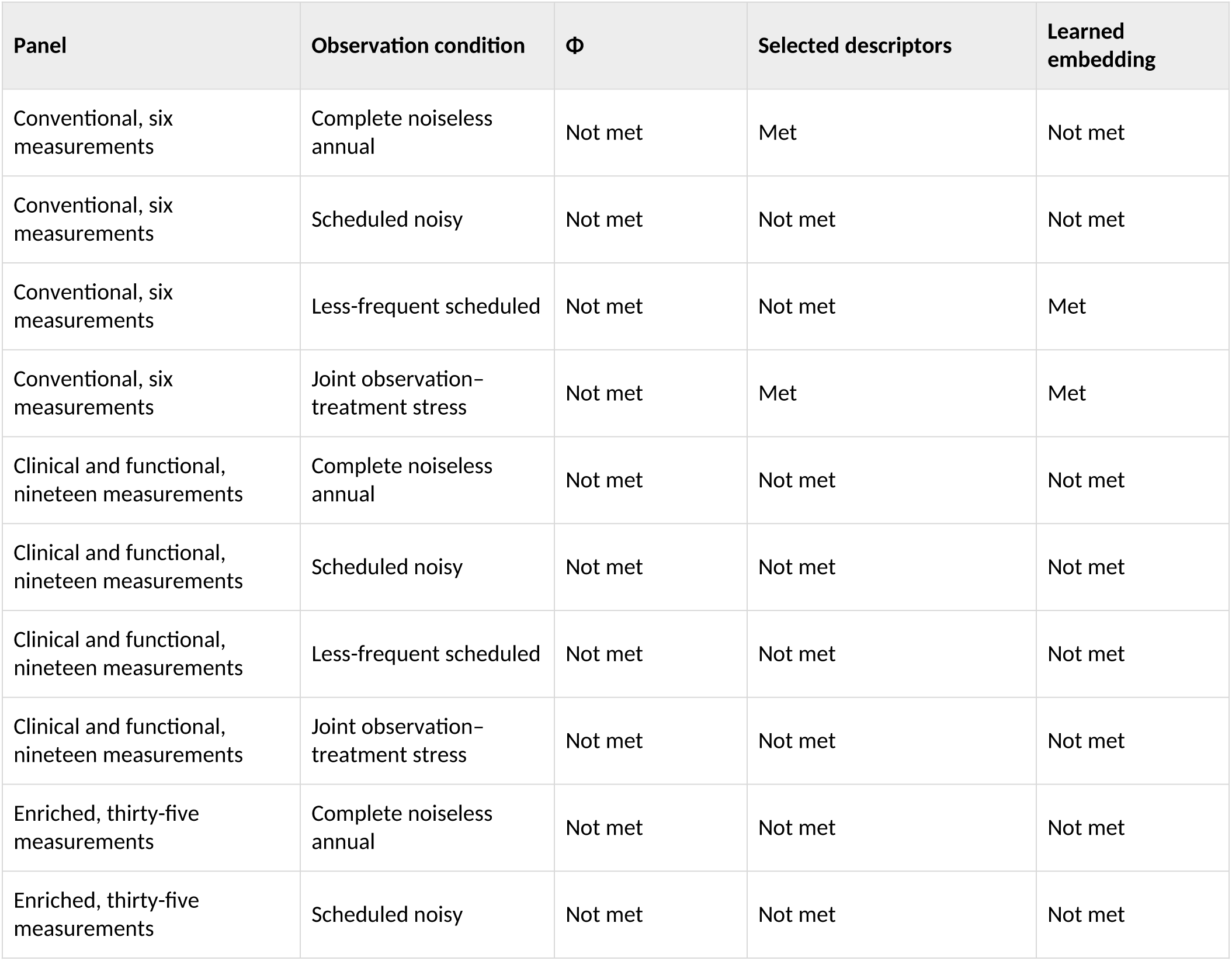
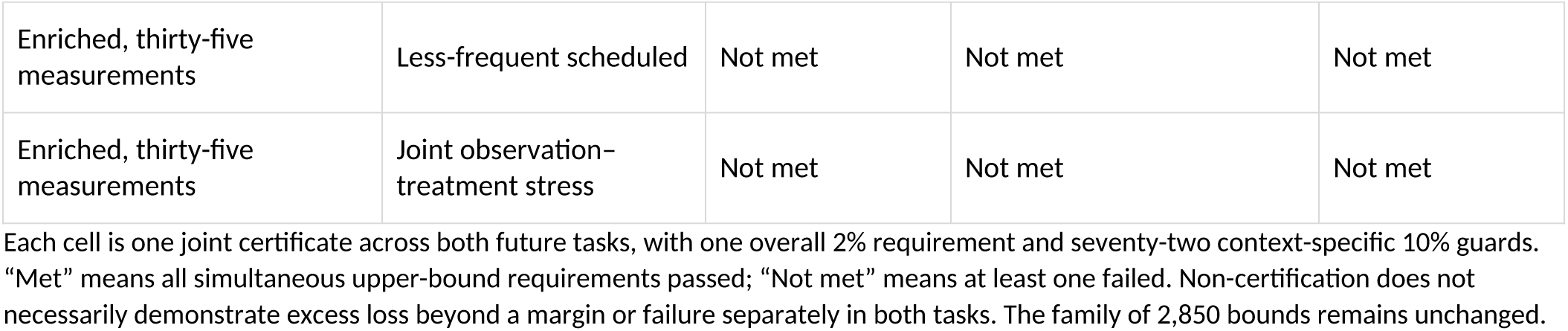
All thirty-six common-representation adequacy decisions.

**Table 3.** All six specialist adequacy decisions under scheduled noisy observation.

| Panel and assigned task | Overall mean relative excess | Simultaneous overall upper bound | Failed guards, out of 36 | Joint requirement |
| --- | --- | --- | --- | --- |
| Conventional; continuation | +0.349% | +1.123% | 2 | Not met |
| Conventional; fixed low load | +1.044% | +3.775% | 4 | Not met |
| Nineteen-measurement; continuation | −4.392% | −2.358% | 3 | Not met |
| Nineteen-measurement; fixed low load | +0.963% | +1.454% | 0 | Met |
| Enriched; continuation | −2.592% | −0.355% | 9 | Not met |
| Enriched; fixed low load | +1.179% | +2.167% | 1 | Not met |

Relative excess refers to predictive loss against the corresponding fitted available-history reference, not information lost or individuals affected. The overall upper bound must be at most 2%; every one of thirty-six guards must be at most 10%. “Fixed low load” replaces the historical withdrawal label without changing the task. The setting assigns controlled inputs to 0.05 and does not guarantee a reduction for every individual.

The reasons for non-certification differed. Under complete noiseless conventional observation, selected descriptors had mean relative excess loss of −1.029%, a simultaneous overall upper bound of −0.688%, and all seventy-two guards passed. Under scheduled noisy observation, their overall estimate was +0.817%, within the 2% allowance, but the upper bound was +2.868% and eleven guards failed. The learned embedding’s scheduled-noisy overall upper bound was +1.262%, but thirteen guards failed. Under less-frequent observation its overall upper bound was +0.481%, and all guards passed.

Among the thirty-seven non-certified groups, eighteen failed overall and guard requirements, eighteen passed overall but failed guards, and one failed only overall. Across 393 failed predicates, 321 had point estimates within the relevant margin but upper bounds above it. Other failures included substantial adverse estimates. Under noiseless conventional observation, all six failed Φ guards concerned nonlinear fixed-low-load prediction, including waist relative excess +23.119% with upper bound +29.464%, and glucose +17.873% with upper bound +22.636%. Uncertainty-limited certification and sizeable estimated excess therefore both occurred.

Only the nineteen-measurement fixed-low-load specialist certified, with mean relative excess +0.963%, overall upper bound +1.454%, and no failed guards. Its continuation counterpart had a mean relative excess loss of −4.392% and upper bound −2.358%, yet three guards failed. The certified specialist reused the common selected-descriptor recipe; that recipe’s two-task certificate failed because its nine failed guards all concerned continuation. Thus, failure of the joint certificate did not invalidate the task-specific certificate. Comparisons between observation settings also used different available-history references and did not establish that a sparser schedule improved absolute prediction.

### Small outputs could require extensive inputs

The conventional-panel learned embedding displayed eight coordinates, depended on 72 upstream engineered value features and had a nominal input-field count of 117 (Figure 2; Table 4). Conventional Φ also displayed eight values, but its nominal input-field count was 53. Selected descriptors retained 48 values with a nominal input-field count of 133. All three conventional value constructions drew on six measurement channels, while their temporal requirements and auxiliary inputs differed.

**Figure 2.**
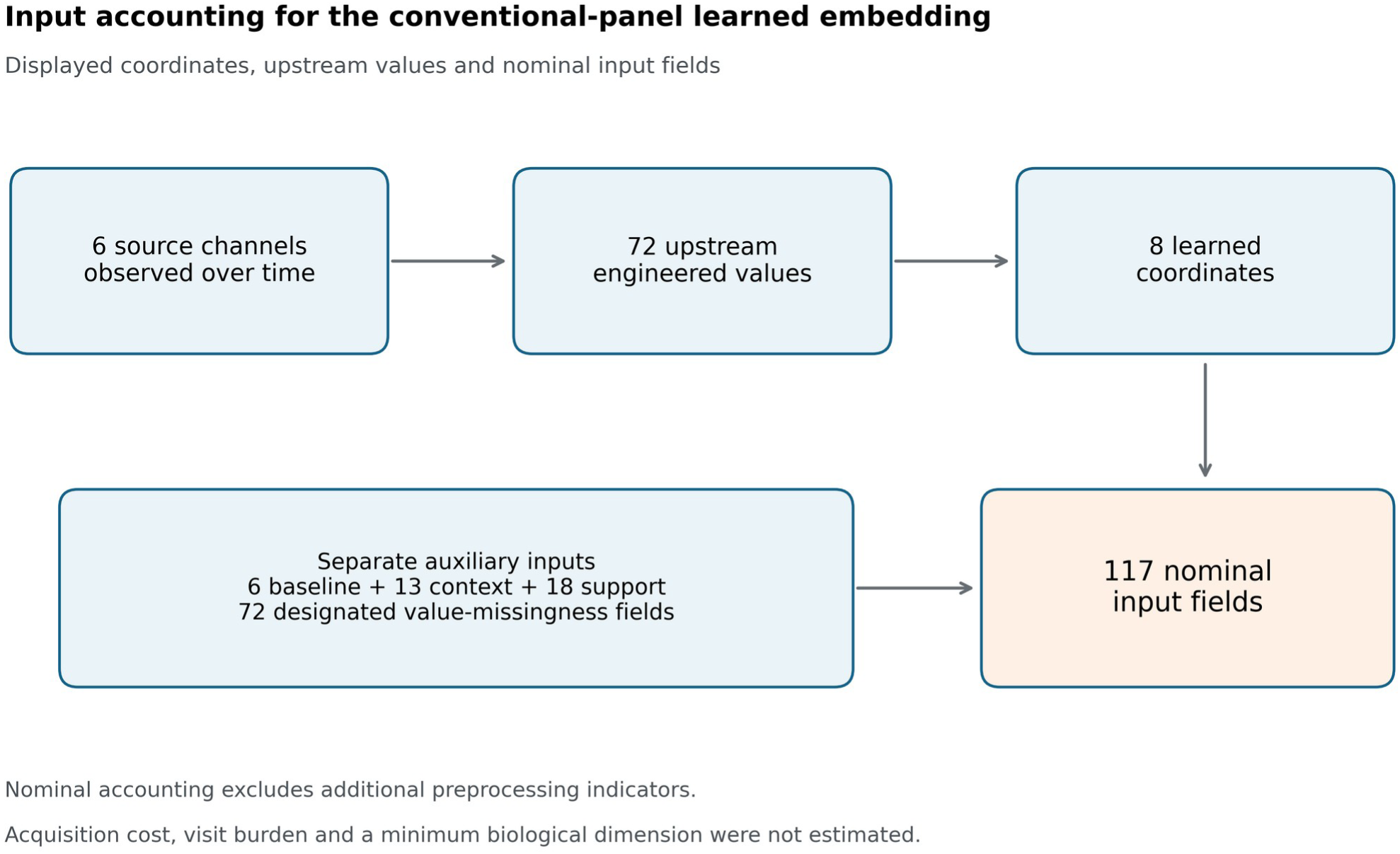
Eight displayed coordinates depended on a larger observation-derived input construction. The conventional-panel embedding used 72 upstream engineered values from six measurement channels; the stated accounting convention gives 117 nominal input fields after adding baseline, context, support and designated value-missingness fields. Additional indicators introduced by downstream preprocessing are excluded. Coordinates, features, input fields and measurement channels are different quantities. The diagram does not estimate cost, visit burden or a minimum biological dimension.

**Table 4.** Displayed values, upstream features and nominal input-field counts.

| Panel | Representation | Upstream value features | Projection coordinates | Nominal input-field count | Channels in value block |
| --- | --- | --- | --- | --- | --- |
| Conventional | $\Phi$ | 8 | — | 53 | 6 |
| Conventional | Selected descriptors | 48 | — | 133 | 6 |
| Conventional | Learned embedding | 72 | 8 | 117 | 6 |
| Nineteen-measurement | $\Phi$ | 16 | — | 121 | 10 |
| Nineteen-measurement | Selected descriptors | 215 | — | 519 | 19 |
| Nineteen-measurement | Learned embedding | 215 | 12 | 316 | 19 |
| Enriched | $\Phi$ | 32 | — | 217 | 14 |
| Enriched | Selected descriptors | 311 | — | 775 | 35 |
| Enriched | Learned embedding | 311 | 12 | 476 | 35 |

Nominal input-field counts comprise representation values or learned coordinates, their designated value-missingness fields, and raw baseline, context and support fields under the stated accounting convention. They exclude additional indicators introduced by downstream preprocessing and do not report measured post-preprocessing matrix widths. Unsupported-column omission and constant-column handling depend on the fitted population.
The last column concerns distinct channels used by the selected value block. For the larger-panel $\Phi$ blocks, counts of ten and fourteen are not verified total acquisition unions once appended baseline, context, support and missingness fields are considered. Nominal input-field counts follow the stated accounting convention and are not assay counts. A dash indicates no learned projection. Actual usable nonconstant columns can differ by cohort.

For the nineteen- and thirty-five-measurement panels, the learned embedding retained twelve coordinates from 215 and 311 upstream values, with nominal input-field counts of 316 and 476. The corresponding selected-descriptor constructions retained the entire candidate pools, with nominal input-field counts of 519 and 775; selection did not necessarily reduce dimension. Φ retained sixteen and thirty-two values, with nominal input-field counts of 121 and 217. Its reported ten- and fourteen-channel counts describe the selected value blocks, not a verified union across the complete predictor inputs.

Continuation specialists displayed twelve values with nominal input-field counts of 61, 113 and 177 across panels. None of these counts was an assay count. Nor did a shared channel imply a shared measurement burden: a current reading and a ten-year descriptor could require different visits and coverage. The accounting supported a distinction between apparent compactness and upstream dependence, but did not establish reduced acquisition, lower cost or clinical feasibility for a particular recipe.

### Repeatability and predictive accuracy did not coincide

In current-state-sufficient continuation diastolic-pressure prediction, Φ-derived hard clusters had lower remeasurement spread than Φ: 0.00082159 versus 0.00936694 on the frozen squared scale (Table 5). Their clean-current-input displacement was also lower, 0.00049795 versus 0.00438716. Nevertheless, Φ had lower error against the noisy future measurement, 0.35169670 versus 0.36863765. Against the noiseless realised future reference, the ordering reversed: 0.04457388 for clusters versus 0.06622806 for Φ. The predictions and fitted models were identical across the two scoring comparisons.

**Table 5.** Repeatability and three reference-dependent prediction endpoints.

| Endpoint | $\Phi$ | Φ-derived hard clusters |
| --- | --- | --- |
| Prediction spread across observation replicas | 0.00936694 | 0.00082159 |
| Mean squared error against noisy future measurement | 0.35169670 | 0.36863765 |
| Mean squared error against noiseless future reference | 0.06622806 | 0.04457388 |
| Squared displacement from the clean-current-input prediction | 0.00438716 | 0.00049795 |

Current-state-sufficient continuation diastolic pressure; conventional panel; scheduled noisy observation; Ridge regression. Values are equal-cohort means across twenty cohorts, with thirty replicas for each of 256 held-out individuals per cohort. All endpoints use the frozen training-target squared scale and are dimensionless. No new significance tests are attached. The noiseless future is one realised biological trajectory. Clean-current-input displacement uses a different anchor from future error.

The noiseless-error cohort ranges were 0.054676–0.091213 for Φ and 0.035572–0.054778 for its clusters. These were observed ranges, not confidence intervals. In nonlinear fixed-low-load fasting-glucose prediction, clusters again had lower spread, 0.0086349 versus 0.0115468, but Φ had lower error against both noisy and noiseless futures: 0.145715 versus 0.175484, and 0.096137 versus 0.114567, respectively (Figure 3). Current Measurements had lower spread, clean-input displacement and both errors than their own clusters across the nonlinear twelve task-channel combinations. No universal relation between stability and accuracy emerged.

**Figure 3.**
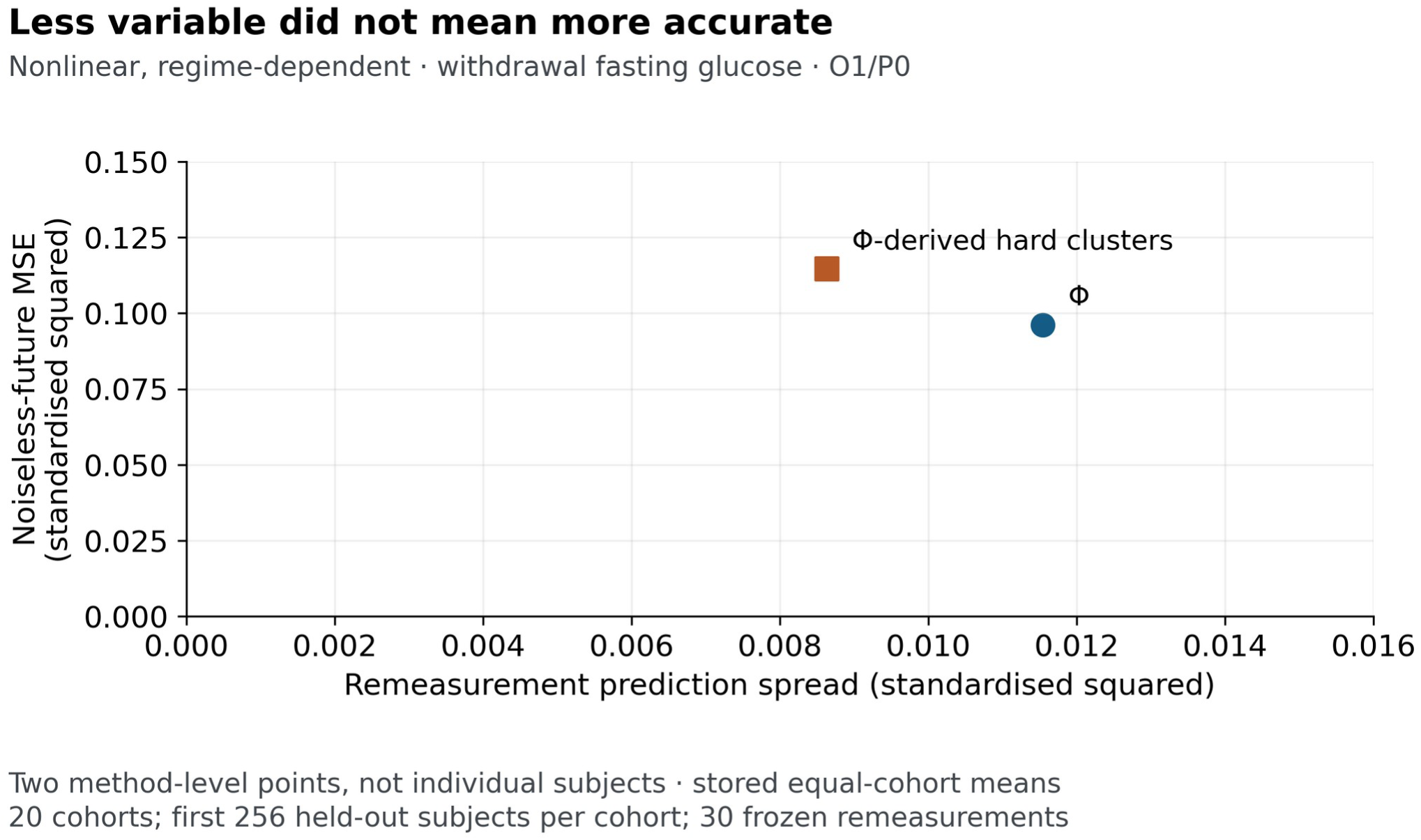
Greater repeatability did not ensure greater accuracy in the nonlinear fixed-low-load example. Φ-derived clusters had lower prediction spread but higher noiseless-future error than Φ for fasting glucose. Points are descriptive equal-cohort means across twenty cohorts, with thirty remeasurements for 256 held-out individuals per cohort. Both axes use the frozen channel-specific squared scale. The noiseless future is one realised trajectory. Any historical “withdrawal” label denotes the branch assigning inputs to 0.05, not guaranteed input reduction. O1/P0 denotes scheduled noisy observation of the conventional six-measurement panel.

Label-matching comparisons also required care. In nonlinear continuation waist-cluster prediction, displacement from the original saved prediction was 0.00016193 among matching-label replicas and 0.71671247 among differing-label replicas. An adaptive-reference continuation HDL example showed the reverse ordering, 0.01117586 versus 0.01104786. The same individual could contribute to both strata through different replicas, so these were conditional descriptive summaries, not causal effects of changing a label.

Pipeline reconstruction showed exact preservation of raw current values by the six current Φ coordinates. All 245,760 evaluable learned-embedding subject–coordinate pairs varied across replicas. Where defined, 509,721 selected historical descriptor entries and 850,761 historical embedding-native inputs remained invariant, consistent with the fixed-history design. An adverse current-state-sufficient cohort-15 embedding example remained in the record: fixed-low-load systolic-pressure noiseless error was 0.729113 versus 0.124150 for its clusters. These observations located sensitivity; they did not establish that sensitivity alone caused the prediction differences.

### Natural tracking included weak and undefined associations

The coordinate analysis contained 26,880 cohort-specific associations. Of these, 25,440 were defined, including 442 negative values; 1,440 were undefined. Under complete noiseless annual observation, defined correlations ranged from 0.3459 to 0.8786. The corresponding ranges were −0.0453 to 0.6900 under scheduled noisy observation, −0.0459 to 0.6824 under less-frequent observation, and −0.1316 to 0.6064 under joint observation –treatment stress (Table 6).

**Table 6.** Ranges of defined natural-tracking associations by observation condition.

| Observation condition | Lowest defined correlation | Highest defined correlation |
| --- | --- | --- |
| Complete noiseless annual observation | 0.3459 | 0.8786 |
| Scheduled noisy observation | -0.0453 | 0.6900 |
| Less-frequent scheduled observation | -0.0459 | 0.6824 |
| Joint observation-treatment-policy stress | -0.1316 | 0.6064 |

Extrema are over defined coordinate-cohort Spearman correlations between absolute Φ-coordinate movement and the four-domain latent-change magnitude. They are neither confidence intervals nor estimates of a causal noise effect. The stress condition also changes treatment policy. There were 25,440 defined and 1,440 undefined correlations; undefined values are not zero.

Selected examples illustrate the heterogeneity (Figure 4). Liver attenuation under accumulated-burden, noiseless enriched observation had mean cohort correlation 0.8617, with range 0.8491–0.8786. Diastolic pressure under current-state-sufficient joint stress had mean 0.00762, with range −0.05432 to 0.05323. These examples used different coordinates and conditions; their contrast was not an estimate of the causal effect of noise. Negative associations concerned the magnitudes of changes and did not, by themselves, indicate a reversal of physiological direction.

**Figure 4.**
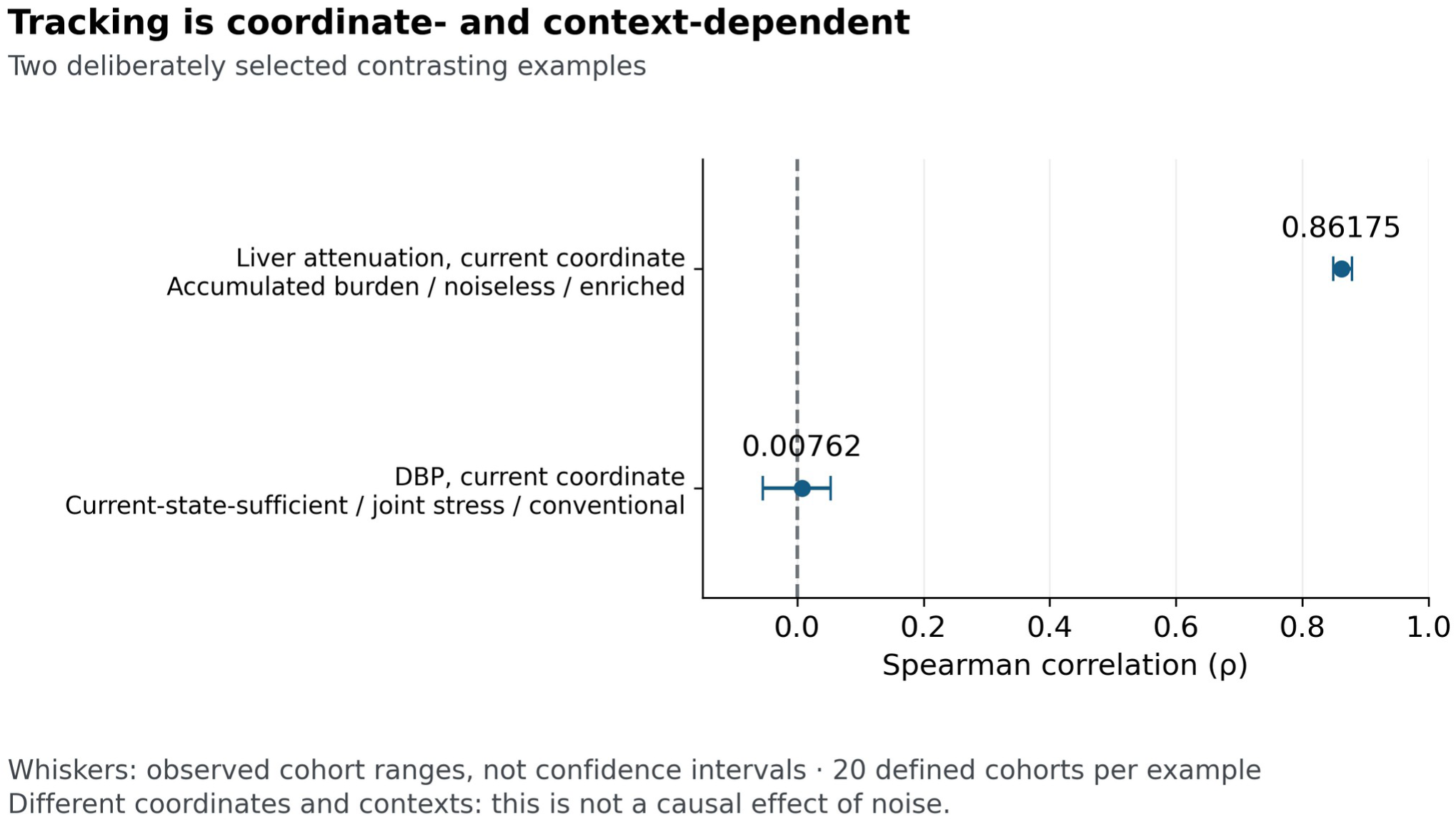
Natural tracking varied across coordinates and contexts. The selected strong and near-zero examples correlate absolute Φ-coordinate change with the four-domain latent-change magnitude. Points show mean cohort Spearman correlation; lines show observed cohort minima and maxima, not confidence intervals. Each example has twenty defined cohorts. The examples use different coordinates and observation conditions and do not isolate a causal effect of measurement noise.

All 50,880 eligible grouped summaries conditional on unchanged Φ-cluster labels had positive mean absolute coordinate movement. This established movement within unchanged labels at the grouped-summary level, not movement in every individual. Across coordinate–subject pairs, 22,760,859 were finite and 4,737,381 unsupported. All 1,440 undefined cohort correlations occurred in the joint stress condition because laboratory-history windows lacked support. Unsupported histories were not interpreted as stable physiology. Whole-representation distances and the unavailable year-10 selected, learned and research-probe comparisons remained unestimated.

### Controlled responsiveness depended on the reference and denominator

The controlled procedure completed all 120 contexts and 30,720 subject–cell identities. All 21,060,288 numerically resolved responses in predeclared directional families agreed with their assigned direction; another 24,485,808 eligible responses were unresolved. This result applied only where a direction had been specified. It did not establish directions for arbitrary learned axes, nor convert unresolved changes into evidence of no effect.

Direction agreement did not ensure monotonic dose ordering. Of 31,457,280 three-dose sequences, 321,398 were classified as mixed. All 42,302 mixed Φ sequences arose in nonlinear dynamics. A further 5,609,712 final-coordinate rows were undefined. These were repeated analytical records, not independent experimental subjects. Numerical tolerances distinguished resolved from unresolved calculations; they were neither statistical significance thresholds nor clinical change thresholds.

In the paired stochastic lane, 40,685,742 of 47,127,708 eligible response rows were smaller than one matched measurement standard deviation, giving 86.331% (Figure 5; Table 7). The remaining 6,441,966 rows exceeded one standard deviation; 6,222,081 of these, or 96.587%, retained their corresponding category. A separate summary found category silence in 61,085,344 of 62,291,277 numerically resolved final-coordinate response rows, or 98.064%. At the representation–category-pair level, 97,191 of 4,423,680 pairs switched category, or 2.197%.

**Figure 5.**
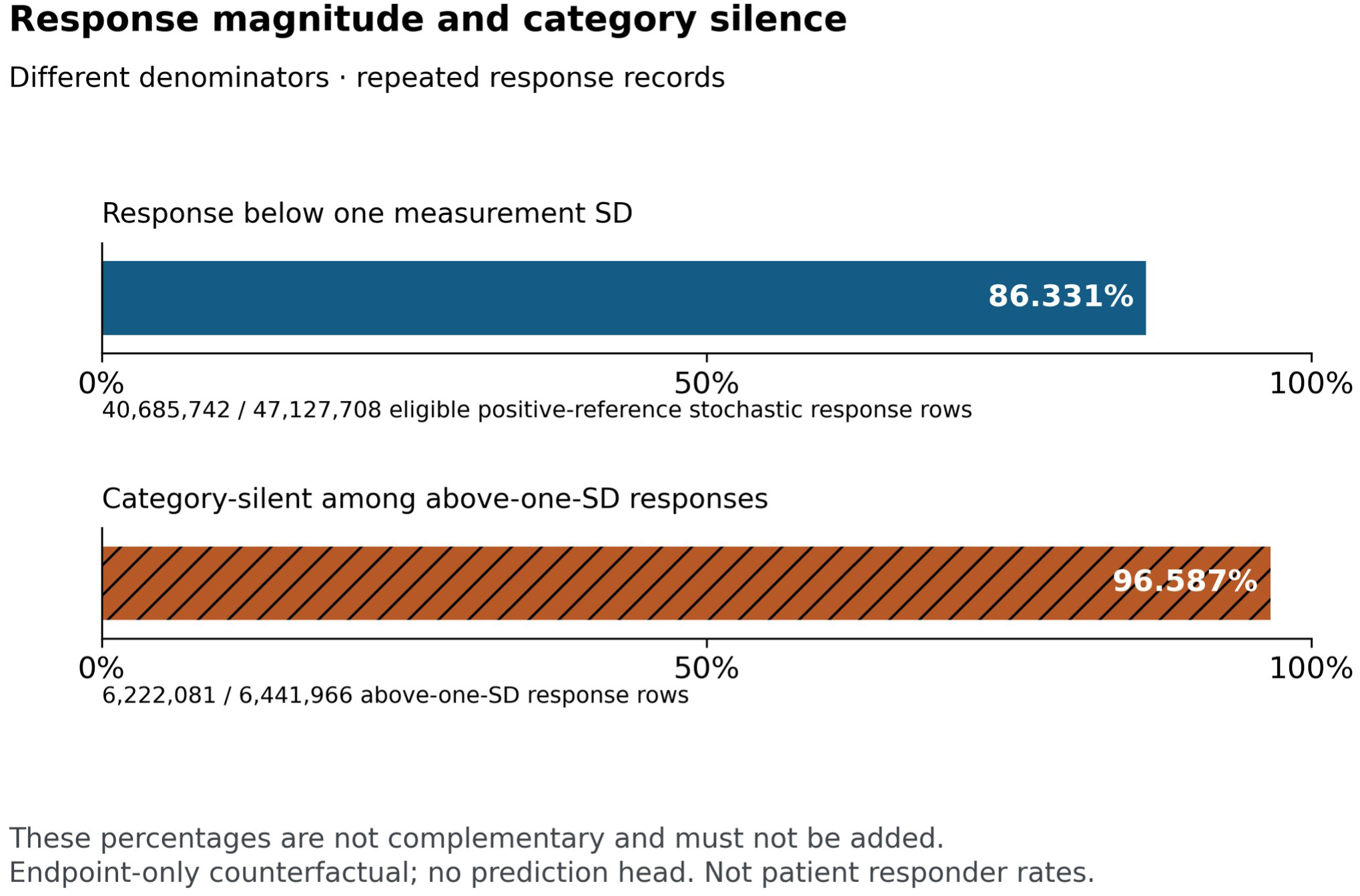
Measurement-relative magnitude and category silence describe different conditional populations. Among 47,127,708 eligible stochastic response records, 86.331% were smaller than one matched measurement standard deviation. Among the 6,441,966 records exceeding that reference, 96.587% retained their corresponding category. The denominators repeat individuals and are not patient counts. These percentages must not be added. Earlier observed history remained fixed, and subsequent prediction loss was not evaluated.

**Table 7.**
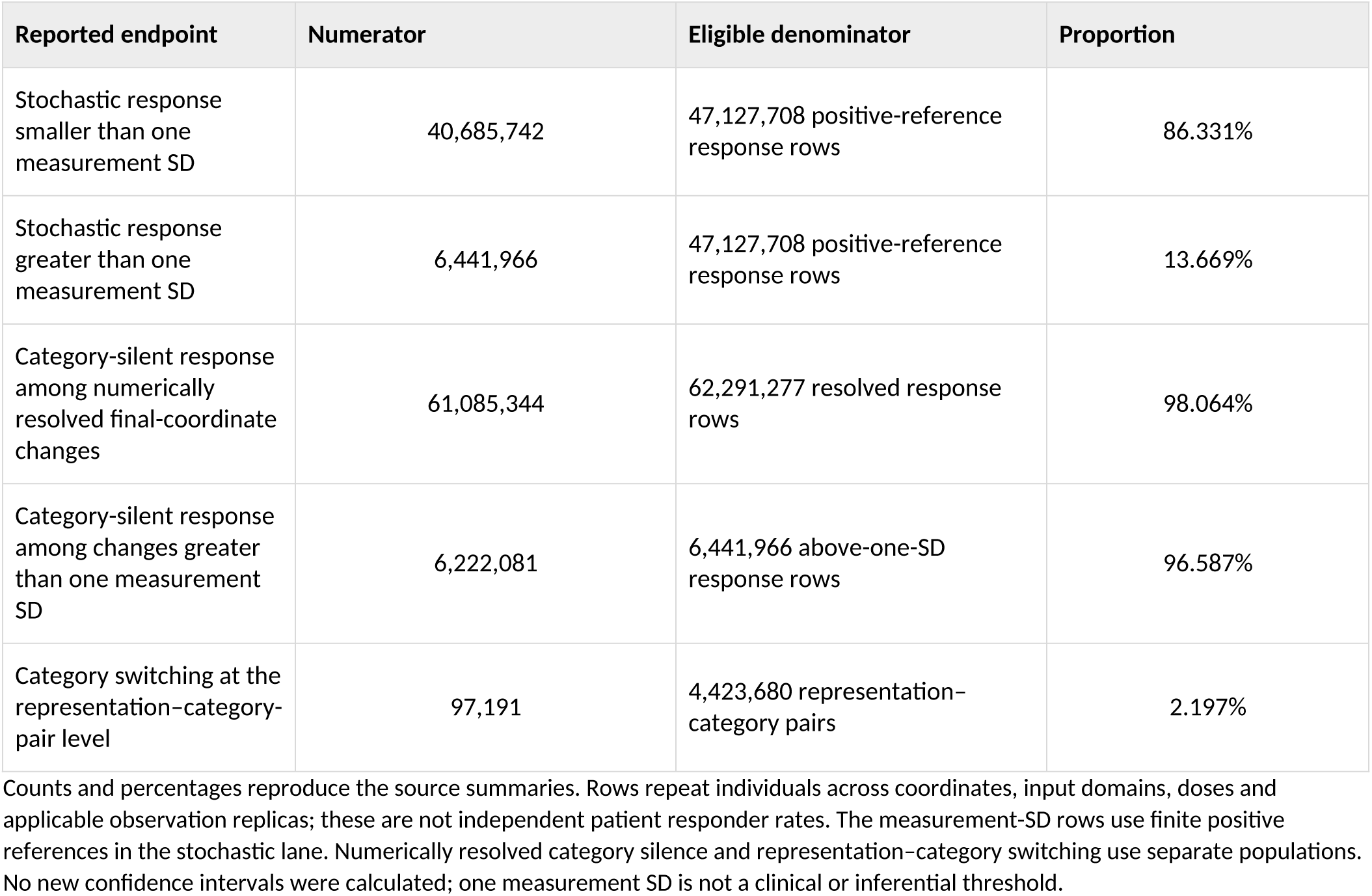
Controlled endpoint response counts with their distinct denominators.

These fractions used different eligibility rules and denominators. They cannot be added, interpreted as independent patient proportions or substituted for equal-cohort summaries. For 16,077 resolved selected-descriptor responses with zero reference variance, R remained undefined. No infinitesimal constant or infinite response was substituted. The findings demonstrated that category silence could accompany responses larger than the chosen measurement reference, while also showing that most eligible stochastic responses were smaller than that reference. Subsequent predictive loss was not measured in this procedure.

## Discussion

### Adequacy is a declared requirement rather than a general rank

The principal finding is that predictive adequacy, compactness, repeatability and responsiveness supplied different evaluations of the same representation strategies. Five joint certificates were obtained, but none of Φ’s twelve common groups certified. This does not establish a universal failure of continuous representation, and it does not justify reading a failed certificate as demonstrated excess loss in every context. The 321 failed predicates with estimates inside their margins show how uncertainty affected certification. The large adverse Φ estimates in nonlinear fixed-low-load prediction show why uncertainty was not the only explanation.

The specialist example makes the scope of a certificate concrete. A recipe met the fixed-low-load requirements even though its two-task conjunction failed. Aggregating tasks into a single verdict therefore changed the question, without changing those task-specific predictions. A certificate also depended on a fitted available-history reference. Changing observation conditions changed the information available to that reference, so certification under a sparser schedule could not establish that sparse measurement was intrinsically better. Formal sufficiency and operational adequacy should remain distinct (Dawid, 1979).

### Information requirements extend beyond the displayed representation

The eight-coordinate embedding illustrates why output size is an incomplete account of burden. Its construction required 72 upstream values and additional fields that gave a nominal input-field count of 117 under the stated accounting convention. These quantities describe different stages. A coordinate count is not an assay count, and a feature count does not identify how many visits are needed to estimate a historical descriptor. The larger selected-descriptor recipes also show that a selection procedure can retain its whole candidate pool.

The present ledger does not establish the full acquisition union for every representation. In particular, the ten- and fourteen-channel counts for Φ value blocks do not establish total ten- or fourteen-assay requirements when appended baseline or support fields may use additional channels. Before a clinical burden comparison, the full dependency chain would need to identify each measurement, occasion, coverage requirement, missingness rule and access constraint. Cost and patient burden would then require their own observations, rather than being inferred from numerical width.

### Movement requires an explicit measurement and physiological reference

Prediction spread measured repeatability under the specified current remeasurement procedure. It did not measure closeness to a future outcome. The same predictions could change ordering when scored against noisy or noiseless realised futures, while a more stable clustered predictor could be less accurate against both. Stability may result from suppressed distinctions as well as reduced measurement sensitivity. These findings reinforce the need to separate the properties of an observation, its representation and its predictive use (Shmueli, 2010).

Natural tracking and controlled response answered further questions. Natural tracking associated absolute coordinate movement with a model-defined latent-change magnitude and retained weak, negative and unsupported cases. The controlled procedure compared matched endpoints under an imposed input change, with earlier observed history held fixed. Neither supplied a universal physiological movement score. Their interpretation is compatible with measurement-invariance concerns in empirical metabolic scoring, where the relation between an indicator and its construct requires examination over time (Barbosa-Leiker et al., 2011).

Most eligible stochastic responses were below one matched remeasurement standard deviation, yet category silence remained frequent among those exceeding it. Neither statement contradicts the other because the second conditions on a different subset. One measurement standard deviation was a descriptive reference, not the standard error of a paired response. Values below it were not proved undetectable; values above it were not proved clinically important. Moreover, the endpoint experiment did not test whether a silent change caused additional future prediction error. That question would require its own outcome evaluation.

### Limits and empirical implications

The findings are conditional on stipulated synthetic laws. Narrow repeated-cohort uncertainty does not quantify uncertainty about whether the generating laws describe human physiology. The four burden domains and historical carriers were chosen model constructs, and the stress observation condition altered treatment policy as well as measurement. The available-history predictor was not an optimal estimator, and recipe selection used a bounded development menu. No result establishes a globally minimal or sufficient human physiological state.

The analyses also differed in populations, chronology and support. Remeasurement and controlled response used 256 held-out individuals per context; natural tracking used the full 1,023. Their overlapping cohorts and repeated records cannot be pooled as independent evidence. Later reconstruction and controlled-response work retained their separate status, three historical predictive tasks remained unavailable, and some whole-vector endpoints could not be evaluated using only information available by year 10. Source checks and bounded replay supported specific computational relationships but did not certify unrestricted reproduction of every historical stage. The disclosed aggregate results permit inspection, but regeneration requires the additional assets listed in Supplementary Methods S7.

An empirical programme should therefore define the question before choosing the representation: which outcome and horizon must be predicted, what excess loss is acceptable, what observations are available, and what kind of change matters? Prospective comparison should measure the entire input burden, use matched information sets where transformation is the question, and retain realistic observation schedules and reporting delays. Repeatability studies should identify their measurement reference; responsiveness studies should connect movement to outcomes that matter for the intended use. Established severity scores and longitudinal models provide appropriate comparators rather than being displaced by a claim that a dynamical formulation alone guarantees improvement (Gurka et al., 2014; Breeze et al., 2016; Vishnu et al., 2015).

## Conclusions

In these controlled synthetic systems, meeting a predictive requirement, displaying few coordinates, repeating stably and responding to changing input were distinct achievements. Five of forty-two adequacy groups certified, while input accounting and movement analyses exposed additional dependencies and limitations. An informative evaluation preserved task scope, full input requirements, measurement references, adverse findings and unsupported cases. The results support that evaluation strategy; they do not establish a universally preferred representation, human physiological validity or therapeutic benefit.

## Supporting information

Shared study methods for the METLAS companion papers

## Declarations

### Relationship to the companion article

This article and the companion manuscript “Predictive consequences of alternative representations of metabolic measurements” report distinct questions from the same synthetic study programme. They share generating and observation methods and overlapping simulated cohorts; they are not independent replications. The shared Supplementary Methods document specifies the bounded study design.

### Ethics and consent

The study used computationally generated synthetic cohorts. It involved no recruitment of or interaction with human participants, no patient or participant records, no identifiable personal information or biological specimens, and no animals. No ethics committee approval or formal exemption was obtained. Informed consent was not applicable.

### Data and code availability

The article and shared Supplementary Methods disclose the bounded synthetic model, observation mappings, locked Φ coordinates, representation and fitting rules, and the displayed result tables. The broader METLAS framework, confidential calibration resources and production implementation are outside the reported study and are not distributed.

The accompanying materials do not include the executable study implementation, complete selected-feature and embedding-input rosters, full-precision constructed-query records, complete machine-readable result families or paired-cohort loss arrays. Their absence limits exact computational reproduction. Requests for study-specific supporting materials may be directed to the corresponding authors and will be considered subject to ownership and release permissions; access is not guaranteed. Supplementary Methods S7 describes the materials provided and the remaining reproducibility limitations.

### Author contributions

Dr Mwangi Njenga: Conceptualization, Methodology, Formal analysis, Investigation, Software, Writing, Original draft

Dr Waweru Peter: Conceptualization, Supervision, Writing-Review and Editing, Project administration

### Use of AI assistance

AI-assisted tools were used in computational implementation and analysis, and in the organisation, drafting and editing of the manuscripts. The authors are responsible for the scientific content, interpretation and final text.

